# Self-organized patterning and morphogenesis in embryonic epithelia from antagonistic signals and fiber networks

**DOI:** 10.1101/023069

**Authors:** F.W. Cummings, Kai Lu

## Abstract

A number of universals can be observed in the developing embryos of all phyla. An attempt is made here to describe some of these with a simple model, one consisting of two mutually repelling regions of gene patterning produced by signaling pathways, two acting at each growth phase. The diffusion of ligands is short range, nearest or near neighbors, but the transcription patterns extend over many cellular diameters. The universals discussed are: gastrulation, formation of a blastopore, patterning of stem cells as surrounding compartments (propagating anew with each growth phase, to the adult), the origin of bilaterality, the prevalence of segmentation, and the general ability to regenerate and duplicate. The origin of organ sizes are determined by the parameters of the signaling pathways involved, independent of cell sizes or numbers. The important fiber mesh, or fiber network that can also extend over many cell diameters is also briefly discussed, and is seen as a partner with the signaling pathways in the overall patterning.

## 1) Introduction

The term ‘universal’ is used as meaning ‘common to all animals’, only to convey the sense that all animals exhibit most of these events discussed here. All animals gastrulate, most show periodicities, and the three-layered animals (triploblastic) form the mesoderm from the blastopore ring near the original gastrula opening. Bilateral animals are the most common, and establishing the dorsal/ventral specification comes with the emergence of the mesoderm. Stem cells are present from cnidaria to mammals, and are continually renewed at each growth phase; the question of how and where they are patterned is central. Animals show the ability to regenerate and duplicate, this apparently dependent on the presence of appropriate stem cells (Gilbert 2004; Bryant 1975); the degree of regenerative ability seems to decline with the complexity of the animal, with a large relative deficit seen in mammals.

Our inquiry is to what extent these developmental events can be correlated with one simple, testable model. The relevant ligands of the signaling pathways (SP) are here thought to travel only to nearest neighbors, but give patterns extending over many cell diameters.

It has become increasingly understood how conserved cellular and developmental mechanisms play a major role in changes of anatomy and physiology. While zebras and sparrows look the way they do ultimately because of the genes they carry, the recent ‘Evo-Devo’ revolution has shed amazing new light on development (e.g., Carroll, Grenier and Weatherbee 2005; Muller 2007; Gerhardt and Kirschner 1997). Eukaryotes have by virtue of formation of epithelia and matrix molecules made the crucial transition to numerous cell types and functions. Earthworms and mammals share a surprising number of conserved developmental homeobox genes. There are far reaching consequences for a few key properties of multicellular forms, in particular via combinatorially used switches and ‘tool kit’ genes, and cis regulatory genes. The concept of transcriptional gene regulatory networks is proving to be powerful in this context.

Very important among these developmental effectors are intercellular signaling events consisting of both positive and negative regulators, which are known to be characteristic of a surprisingly wide variety of multicellular processes and developmental events. A potential now exists for better understanding the mechanisms underlying the origin of body plans. Still, a vast number of mysteries of course remain about the unimaginably complex process of animal development, the complexity growing with evolutionary time since the Pre-Cambrian.

A model of gene patterning is proposed in the following, one that involves the effect of the nearest neighbors on a cell, or on a small number of cells, thus introducing cell-cell interactions and signaling pathways. **Each cell** is the potential organizer, or initiator of pattern. An extended model (Section 3) proposes a simple coupling of the gene patterning model to epithelial shape, the latter an essential element in ‘morpho-genesis’.

The epithelial shape is here described at each point (or small region on the middle surface, of the thick epithelial sheet, by the usual Mean (H) and Gauss (K) curvatures (Section 3). The middle surface of a small section may involve four to nine cells. The coupling of the surface curvatures to the gene pattern will be explored.

Of particular interest will be the questions of formation of the blastopore, the ring of tissue just inside the gastrula opening. It is this region from which the mesoderm emerges, and the region of expression of the conserved Brachyury gene (Technau 2001). The mesoderm emerges on opposite sides of this blastopore region (Gilbert 2004, p. 30). The specification of the position of these two points of mesoderm emergence are proposed.

A second focus of the present work is on patterning of stem cells, at each assumed growth phase. Presumably stem cells are not distributed at random, so the question is ‘how are they patterned’, or how are their space positions specified? This question has been explored extensively (e.g., PLOS). If a newt’s tail is excised, the ‘newt-tail’ stem cells act to regenerate the tail; and the excised piece will duplicate when cultured; regeneration proceeds by the local formation of a blastema, a growth zone of mesenchymal stem cells on the stump. This dual property of regeneration/duplication has been studied in detail in the case of imaginal discs (e.g., Bryant 1975). Stem cells are apparently created anew during development with each growth phase. (A ‘growth phase’ will become more clearly defined in the following section).

A new “growth phase”, usually involving cell proliferation, is begun after each steady state is reached; in each growth phase, two (new in general) SP’s are regionally specified, and the regions of developing ‘co-t’ (co-transcription) factors increasingly avoid each other as time advances. A co-t factor specifies a cellular density of factors that enter the nucleus and give rise to changes in DNA transcription. (A common example is β-catenin in the case of the ubiquitous Wnt SP., such A new boundary separating the two cellular regions, consisting of many cells, of two ‘co-t’ repelling regions, emerges as the next steady state approaches, such steady state presaging the beginning of the next growth phase, generally after a brief pause. In cells for which the region between the two steady state co-t factors is below a threshold, the ‘M’ (margin) cells are a boundary region not transcribed, no new genes are activated or deactivated, and thus the region can be considered a ‘stem cell’ region for this growth stage. (Figure 5 is the model pattern shown on a sphere. The stem cell region is an angular region, colored blue in the figure, and lying above the lower threshold of gene activation denoted by the horizontal line **‘t_1_’**).

The five patterning events addressed here are: 1) gastrulation, 2) the blastopore; 3) stem cells; 4) bilaterality and 5) position of outgrowths. The crucially important specification of the **position** of outgrowths (e.g., the ‘**where’** of limbs and antennae, apart from their patterning) from the animal body is proposed, such specification stressing the necessity of simultaneously specifying the anterior/posterior and dorsal/ventral coordinates of such outgrowths.

One motivation for consideration of signalling as a source of patterning is that one species of unicellular choanoflagellates are the closest known relatives of metazoans (Abedin and King, 2008; Nichols, Dagel and King 2009). To discover potential molecular mechanisms underlying the evolution of metazoan multicellularity, the genome of a unicellular choanoflagellate was sequenced and analyzed. The genome contains a large number of genes that encode cell adhesion and signalling protein domains that are otherwise restricted to metazoans. The suggestion is that the adhesion and signaling molecules of the choanoflagellate were precursors of the unicellular to metazoan transition. It prompts one to ask how complex the extra steps might be for the transition, one requiring a separation into distinct gene activation regions of the epithelial sheet of a multicellular animal, including stem cell regions.

Metazoan signaling pathways apparently emerged from these choanoflagellate beginnings. The present model shows that metazoan patterning can be a simple outgrowth of the sequential interaction of two different signaling pathways, the second (Hedgehog, most likely) evolved from the first, and so on (Clevers and Nusse, 2012; Nusse 2003). The Wnt signal transduction cascade (e.g.) controls very many biological phenomena throughout development and into adult life of all animals. Also, aberrant Wnt and Hedgehog signaling, for just one example, underlies a wide range of pathologies and diseases in animals (Taipale and Beachy 2001), including mammals and humans.

Another motivation for the present pattern algorithm of Section 2 are the **odd-leg pairs** of the over 3,000 species of centipede, presumably not an adaptation. This fact is a clue of a pattern mechanism at work. This is a natural outcome of the present work, due to the ‘double-segment’ nature of two co-t factors that avoid each other.

## 2) The Interacting Signaling Pathway model

We explore the question of finding a possible ‘universal’ developmental mechanism(s), one providing distinct and unique specification of different gene transcription regions. This earliest mechanism is no doubt augmented by others as animal complexity evolves beyond the Pre-Cambrian, (but not explored here) e.g., the evolution of neural cells and their filamentous precursors.

This search is motivated by several observations. In spite of numerous extinctions, there are a handful, five or six signaling pathways employed in embryological development, e.g., Wnt, Hh, EGF, Notch, BMP, TGF-β. Another important observation is the conserved nature of the eight HOX genes, and other ‘tool kit’ genes of recent discoveries of ‘Evo Devo’ (Carroll, Grenier and Weatherbee 2005; Carroll 2005; Arthur 2002; Muller 2007). The HOX genes and cis regulatory elements (cis) allow rats, elephants and humans to have about the same number of genes, their HOX and cis elements mostly determinant of their large phenotypic differences. Segmentation is also of interest in this regard because it has apparently evolved repeatedly in a number of animal phyla. There are three distinct lineages of bilaterian animals, each containing one of the three principal segmented phyla. These three are the annelids of the lophotrochozoans, the arthropods within the ecdysozoans, and the vertebrate chordates of the deuterostomes. This seems to reflect a common evolutionary origin of segmentation; the common ancestor was likely a segmented animal (Peel, Chapman and Akam 2005. The prevalence of repeated elements in animals and plants is certainly impressive. The intriguing presence of the number of odd leg pairs of all of the over 3,000 centipede species (Chipman, Arthur and Akam 2004) provides clues as to patterning in a number of animals. The present model provides for such ‘odd leg pairs’ from the outset.

An emphasis here is on pattern formation resulting from only nearest neighbor diffusion, but nevertheless giving gene patterns extending over many cell diameters. The origin of the blastopore and dorsal/ventral patterning, as well as stem cell patterning are a primary focus, with outgrowth placements(e.g., limbs and antennae) a further prediction.

The model envisions a number of transmembrane receptors imbedded in each cell plasma membrane, and each of the different receptor types is activated only by its specific ligand. Two different types of receptors, along with their specific ligands, are constituents of two different interacting signalling pathways at each growth cycle. The ligands and their receptors are highly attracted to each other, so that diffusion of ligand is seen as limited to nearest (or nearby) neighbors.

Wnt is the most thoroughly studied and ubiquitous (Logan and Nusse 2004; Reya and Clevers, 2005; Nusse, 2003; Willert et al. 2003) and the most evolutionarily ancient (Abedin and King 2008) (and used here most frequently for illustrative purposes). Its mutation is commonly associated with human diseases (Thorne et al. 2010). Another very common SP is Hedgehog (Hh) (di Magliano & Hebrok 2003).

The Hedgehog signalling pathway is essential for numerous processes during embryonic development (Ingham and McMahon 2001). Members of this family of secreted proteins control cell proliferation, differentiation and tissue patterning. The Hh pathway remains active in some adult tissues, including control of adult stem cells in the brain and skin. There is also evidence (Taipale and Beachy 2001) that uncontrolled activation of the pathway results in specific types of cancer. Each of the two SPs mentioned above control (at least) two intercellular functions, transcription, and the cytoskeleton (and thus shape).

The pattern model can be stated simply in terms of a cell’s positive and negative feedback actions. The two signaling pathways are chosen generally regionally, but in the case of the first two gastrulation phases of growth, the two signaling pathways likely remain the same, probably Wnt and Hh.

A simple example is shown in Figure 1, where the boundary conditions have been taken to be continuous on both sides and top of a rectangle, similar to those of a (small) torus.

**Figure 1:**
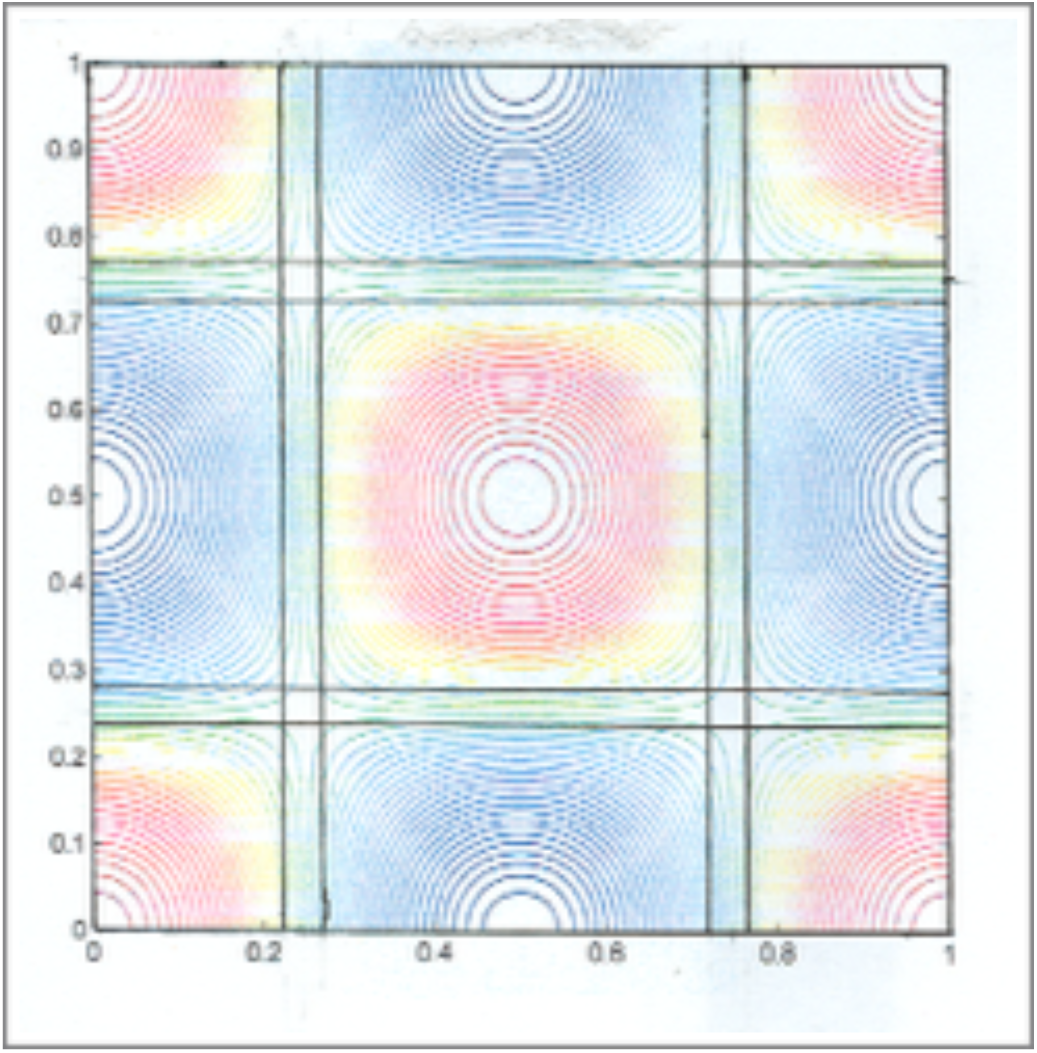
A simple numerical solution of the model is shown on a small torus as an example of the separation of the two co-t factors (shown as red and blue). There will (almost always, depending on the threshold **t_1_)** be a smaller region of ‘non-transcription’ between each co-t region, called the ‘M’ (for ‘middle’) region, where stem cells are patterned for that growth phase. Four regions of overlap or intersection of ‘M’ regions are shown. The ‘torus’ boundary conditions show continuity from top to bottom and left to right.

A few very general properties are assumed for each signaling pathway (SP) in the model: their ligands and receptors are unique to that particular pathway, and each resulting co-transcription factor activates its unique gene network (Wodarz and Nusse 1998; di Magliano and Hebrok 2003)). Ligands are emitted from a cell at a rate proportional to the density of intracellular co-transcription factor (e.g., β- catenin in the case of Wnt, Ci in the case of Hedgehog (Hh), etc.). Emission of the first ligand L_1_ is decreased (negatively regulated) at a rate proportional to the density of co-transcription factor of the second ligand, and vice-versa. Further mathematical details are given in **Appendix A**.

The immense biochemical complexities of the intracellular and extracellular workings of an individual pathway, and the the complex details of interactions of the two pathways, are to a large extent irrelevant for the model. As is well studied in Wnt (Wikramanayake, Huang and Klein 1998), gene activation happens when a certain lower threshold of β-catenin density (β-catenin is one example of a ‘co-t’ factor) is achieved; a single activated surface receptor, or small subset, does **not** lead to gene activation. Here it is assumed that such a lower threshold of co-t is a common property of the several pathways considered. This importantly implies that the nucleus plays a key role in patterning, in setting the threshold levels of gene transcription.

The model consists of a group of cells that have two basic active signaling pathways (SP) interacting at each growth phase. Which two SPs are active is determined by the DNA activation at each cycle in each different region. These SPs act sequentially in pairs, two (possibly different) at each ‘growth phase’.

There are two different corresponding ligand densities in the extracellular space (**labelled L_1_ and L_2_)**, giving rise to two densities (number/area) of co-transcription densities (l**abeled R_1_ and R_2_**).

**Appendix A shows** that the steady sate version of the model is most profitably given in terms of two variables: the ‘Difference’ (‘D’), defined as the difference of the two co-transcription factor densities, i.e., D ≡ R_1_- R_2_, and the Sum ‘S’, where S ≡ K_1_^-2^R_1_ + K_2_^-2^R_2_.

The two resulting very well studied equations, Laplace and Helmholtz, are

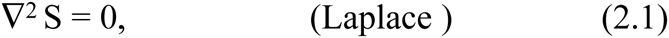

where

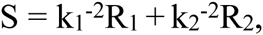

and

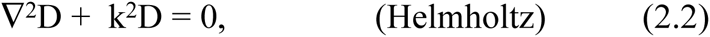

where

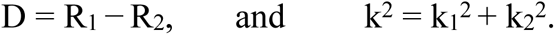

The inverse lengths k_1_ and k_2_ are defined by the parameters of the model by (**Appendix A**)

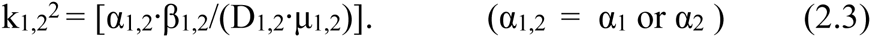

Equations (2.1) and (2.2) are the steady-state equations of the model. Boundary conditions, plus a lower cutoff (assumed the same for both SPs for simplicity) completes the model. These are equations, when coupled to the parameters describing shape, ‘A’, ‘B’ and ‘h’ of the following Section 3, are probed for possible connections to some universals of animal development. From eqs. (2.1) and (2.2), the co-t factors R_1,2_ can easily be reconstructed.

A key ingredient of the model is the fact that all length-scales are determined by the two parameters k_1,2_ of eqs. (2.3); note in particular that there is no mention of individual cells, or cell size. The parameters of the cell signaling pathways determine the organ sizes, e.g., the blastula, which size at the time of gastrulation varies immensely from phyla to phyla and species to species.

The principal and general outcome of the model is that the two co-t regions R_1_ and R_2_ increasingly **avoid each other** as time proceeds, ending at steady state with small overlap of the two co-t regions at the steady state.

The Helmholtz equation (eq. 2.2) produces periodicities in two orthogonal dimensions of the difference function ‘D’, while the Laplace equation produces the maximum smoothness for the sum function ‘S’; Laplace solution maxima and minima always lie on the specified boundary. The Laplace equation is the result of minimizing the squared gradient subject to prescribed values on a boundary, and this latter result is unique.

A simple example of the model on a plane is shown in **Figure 1**, and the two co-t factors R_1_ and R_2_ shown as ‘red’ and ‘blue’. The boundary conditions are here taken as those appropriate to a small torus (or donut), continuous in both directions. Also shown, delineated by straight lines, are the orthogonal ‘M’ regions between the co-t regions, with four ‘M’ region intersections shown. The size or width of M regions are determined by a lower ‘cutoff’, chosen by the **cell nucleus**; below this cutoff, R_1_ nor R_2_ co-t factor density is not sufficient for the co-t factor to enter the nucleus.

There are a vast number of different patterns that can be made by variation of the three parameters k_1_, k_2_ and t_1_. The model of this section is proposed as a base, from which ‘bells and whistles’ can be added (e.g., nonlinearities) if needed.

## 3) Introduction of Geometry of Epithelial Shape

Specification of the geometry or surface shape of a thick surface is essential to any discussion of gastrulation, or development in general (Warmflash, Sorre, Etoc, Siggia and Brivanlou 2014). Previous works have mostly considered a two-dimensional planar geometry, but a recent three dimensional description of epithelial sheets has been given (Hannezo, Prost and Joanny 2014). The specification of any surface involves two radii of curvature (doCarmo 1976) at each vanishingly small (middle surface) region. Since these radii are difficult to connect directly to biochemical and developmental events within the epithelia, we make use of an extant expression that connects the two radii of curvature to three quantities that are most easily accessible to known biological affecters (Cummings 2001, 2006).

The three quantities are the dimensionless apical area ‘A’, the dimensionless basal area ‘B’ and the sheet thickness ‘h’, at each ‘small’ (infinitesimal) middle surface element. The dimensionless quantities ‘A’ and ‘B’ are the apical and basal areas divided by the (square) middle surface element. The three, A, B and h vary across the (imagined) middle surface of the epithelial sheet. Figure 2 shows three different examples of curvatures for (infinitesimal) small sections taken from an epithelial sheet.

**Figure 2:**
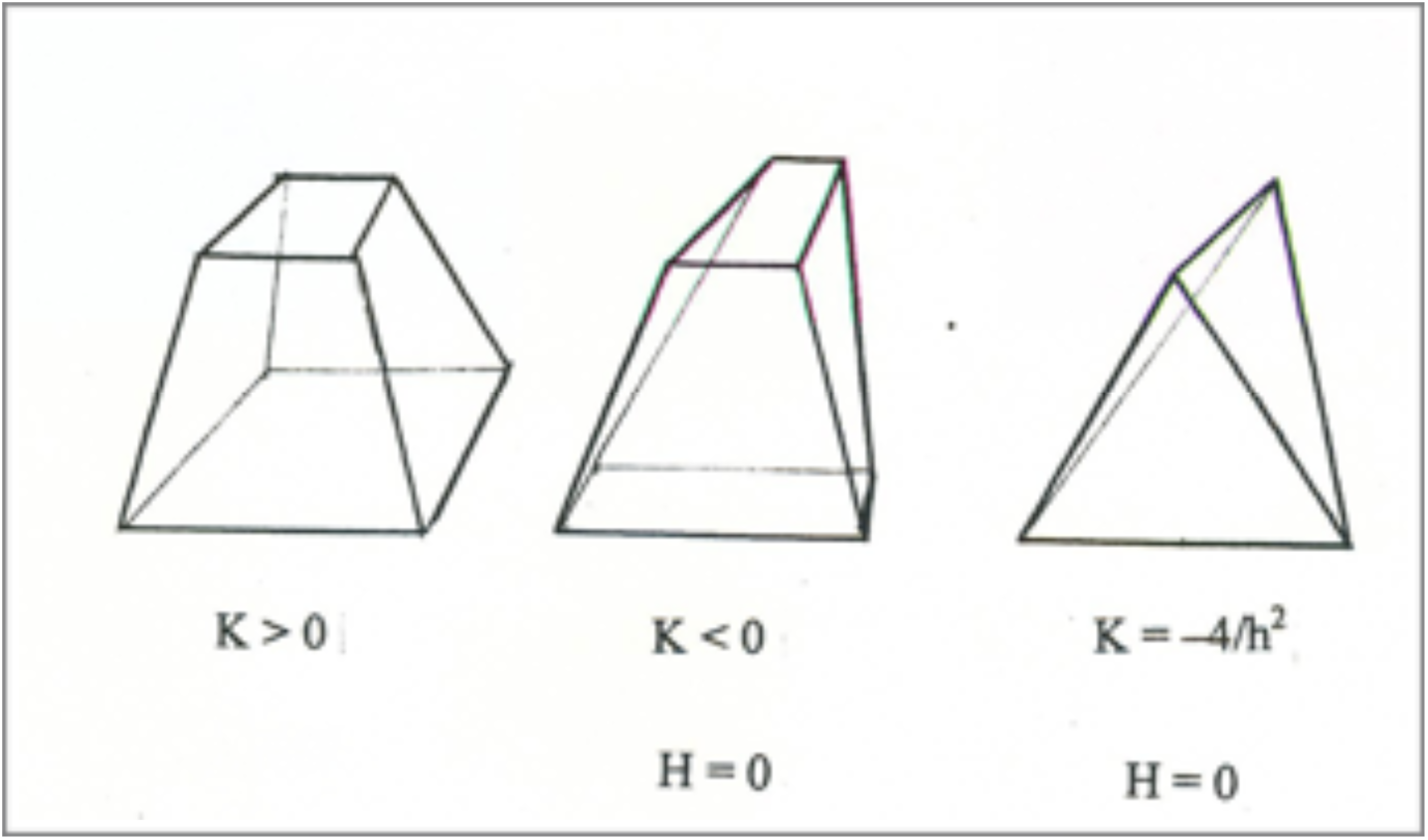
**T**hree examples of different curvature of (∼ infinitesimal in the limit) are shown. The elements are: leftmost (‘a’) shows K > 0, H < 0, A < B, and A, B both as squares; the center picture shows a surface segment with A+B < 2, when K < 0. The dimensionless areas are A = B, and H = 0, K < 0. The right-most extreme ‘saddle’ shape (‘c’) has A = B = 0, and the two limiting case ‘surfaces’ are orthogonal lines, and K = -(2/h) ^2^, H = 0. In all cases the middle surface dividing the apical and basal areas is square. The orthogonal A and B rectangles of ‘b’ and ‘c’ are below the line A+B = 2. The apical to basal height ‘h’ is always described by a straight line through the geometrical center of the three surfaces (A, B and A_m_).

The crucial (invariant) Gauss curvature ‘K’ and the Mean curvature ‘H’ are given exactly by

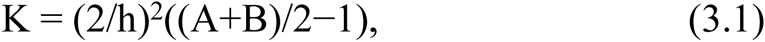

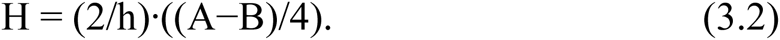

The inequality 4K = (*κ*_1_+*κ*_2_)^2^ – (*κ*_1_–*κ*_2_)^2^ ≤ 4H^2^ serves to restrict most of the A, B plane, and this restriction can be written equivalently simply as

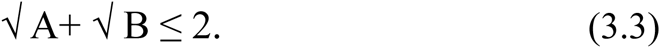

Figure 2 shows three small (infinitesimal in the limit) shapes, with Gauss curvature K > 0 shown on the far left, and two negative K’s on the right, showing the rectangular shapes of A and B in the three cases. The most extreme negative value K = -(2/h)^2^ is shown far right (a collection of such shapes make a ‘saddle’), and with both A and B shown as orthogonal straight lines of zero area, but with a square middle element A_m._

Most shapes are specified by the region of the A, B plane between the line A + B= 2, and the curved line √ A+ √ B = 2; negative Gauss curvature ‘K’ values dip below the A + B = 2, K = 0 line, however; e.g., in the case of gastrulation, near where the two curves meet. Spheres are given when all small elements for a particular A, B and h are given by the equal sign in eq. (3.3), when the sphere radius is R = (h/ 2)/[√ A-1]2, 1 ≤ A ≤ 4; A = 4, B = 0 is a solid sphere, while A = 1 and B = 1 is a sphere of infinite radius. Every small area, apical or basal, is a rectangle, while the middle surface area can be taken **always as a square**. A straight line runs through their three geometrical centers, from apical to basal connecting the three rectangular areas, thus defining an apical to basal direction for each spatial small region.

It is assumed that the shapes (e.g., the examples of Figure 2), are infinitesimal; an infinity of such shapes are required in the limit to constitute a biological surface.

Some limiting shapes are of interest: **a)** The sphere with the most extreme Gauss curvature K is a solid sphere of radius ‘h/2’, when A = 4, B = 0 for an (infinite in the limit) number of such pieces, and K = (2/h)^2^ **; b**) The most extremely negative Gauss curvature surface is a ‘saddle’, formed of shapes with A = B = 0, and K = -(2/h)^2^. This saddle shape is formed from two vanishing areas, each of area A, B = 0, i.e., orthogonal straight lines. However, the area A_m_ of the middle surface of each (infinitesimally vanishing) element is taken as a **square.** including all saddles with H = 0 (and A = B); **c**) A planar small region, K = H = 0, is given by A = B = 1. Any cylinder is a point, or rather an infinity of such points, along the line K = 0, when A + B =2.

Any macroscopic sphere is described by an infinite number of microscopically small shapes, each specified by a single point on the curve of eqn. (3.3), √A + √B = 2. In a similar manner, any cylinder (K = 0) is described by an infinite number of points, all at a single point on the curve A + B = 2.

An unsolved problem is that of closing the system, i.e., one of describing or specifying the three quantities A, B and h as functions of the two co-transcription (co-t) densities, or gene distributions in the surface, at each growth phase. Pattern will then determine shape, and shape in turn will determine gene pattern. Shape determines pattern via the Laplace-Beltrami operator of eqs. (2.1) and (2.2), and Appendix A, since the Laplace-Beltrami operator involves the geometry. The simplest example of such a closing is exhibited in eq. (3.3), and Figure 4.

The example of Figure 4 (below) shows a typical gastrula shape, and the simplest connections have been chosen. This is plausible because the (e.g.) Wnt and Hh pathways both are known to be affecters of the actin cytoskeleton (e.g. Salinas 2007; Sutherland and Witke 1999). There are two (at least) known modes of actions for the Wnt and Hh pathways, and most likely other pathways in embryogenesis also have two or several modes. One most key mode involves collection of cot factors (β-catenin, cis) in the cytoplasm surrounding the nucleus, followed by transcription.

The second mode of Wnt or Hh, and omitted here except for its intracellular action, is apparently for the general purpose of regulating the fiber network. Fibers generally extend out from cell membranes, and create a mesh, a fiber network encompassing the cells of a compartment. The fiber network apparently predates multicellulars, single cells often collecting in masses without genetically distinctive regions, sponges being one of the closest relative to multicellular animals. The three known fibers of the cellular cytoskeleton are most important for controlling animal shape, and of most interest here. The ‘cytoneme’ signaling network (e.g., Roy, Hsiung and Kornberg 2011) introduces a new element beyond the signaling considered so far. Cytonemes are thin tube-like cell extensions capable of conducting signaling ligands over long distances.

Each compartment consists of many cells. Figure 3 shows two compartments on the smallest sphere, the blastula, and four compartments on the two larger spheres. Each compartment is generally suffused by fibers, each cell of a compartment connected to others in the same compartment by fibrous cell extensions, and each compartment often connected in turn to others via a more extensive network.

**Figure 3:**
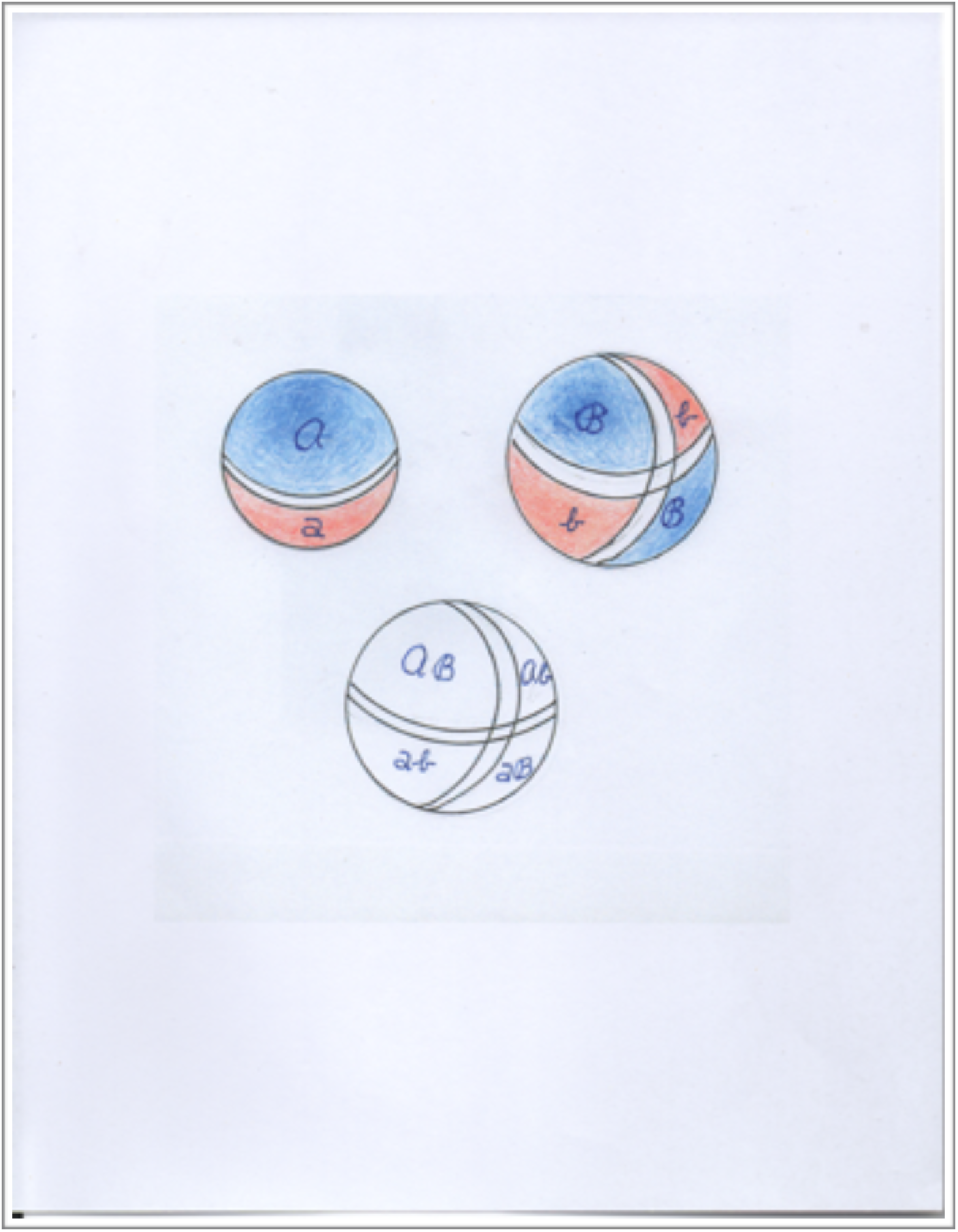
Three pattern functions, solutions of the model, are shown on spheres, where the angle θ runs from pole to pole, i.e., from 0 to π. The smallest sphere (top left) shows two co-t factors, R_1_ and R_2_, with their maxima at opposite poles of the sphere. The two functions decrease toward the equator, and overlap near the equator. The symbol t_1_ marks the lower threshold below which no co-t is allowed to enter the nucleus (c.f., Figure 5) (and is taken as the same for both co-t factors for simplicity). Also indicated by a clear (no color) circle near the equator is the ‘M’ region of ‘no transcription’, and taken to be the ‘stem cell’ region. The smallest sphere of phase one growth has a size of (kR)2 = 2 (cf. Appendix A). The larger sphere (top right and bottom) has an area **three times larger** than the small sphere, with an area of (**kR)^2^ = 6** (Appendix B). Two different co-t factors are shown as ‘ℬ ‘and ‘b’, indicating their four regions of transcription activity. There are now two ‘strips’, specifying the two different ‘M’ regions, with two intersections. The lower sphere indicates the four different, unique specifications of four differential regions, indicated as: *A*ℬ, *A*b, aℬ and ab. The two ‘stem cell’ intersections are proposed as the place of two origins of the mesoderm. (Here ‘A’ denotes ‘script A’)

We present below (eq. (3.3)) the simplest model of the connection between the co-t factors R**_1,2_** and the three geometry factors A, B and h. When one co-t factor, e.g., R**_1_** is small (and transcription of type #1 is ‘off’), A is also relatively small, and vice versa. By the model, R**_1_** small implies R**_2_** is relatively large (Figure 5), and B is then relatively large; the cell has a small ratio of apical to basal area. The R_1,2_ of the model refer to the level of co-t factor accumulating near the nucleus, and shows a gradient pattern extending over many cell diameters (∼ **k^-1^**).

The simplest linear relation, and to be noted, one giving the typical gastrula shape is

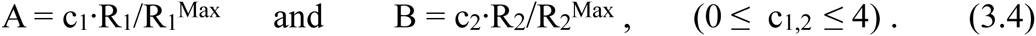

The sheet thickness ‘h’ has been held constant (for simplicity only). Figure 4 in conjunction with Figure 5 is one example of the many possible, but similar shapes, for just one choice of c_1_ and c_2_. In Figure 5, the angle ‘θ’ is zero at the point ‘a’, while the point ‘e’ corresponds to θ = π. When R_1_ (e.g.) is largest, the apical area A is largest (a prediction), and A decreases when R_1_ decreases. The two compartments, described by R**_1,2_** are coordinated, according to the model of Section 2: apical area ‘A’ is then large when transcription (R_1_) is large by eq. (3.3) and vice versa, and while area ‘B’ increases, ‘A’ decreases.

**Figure 4:**
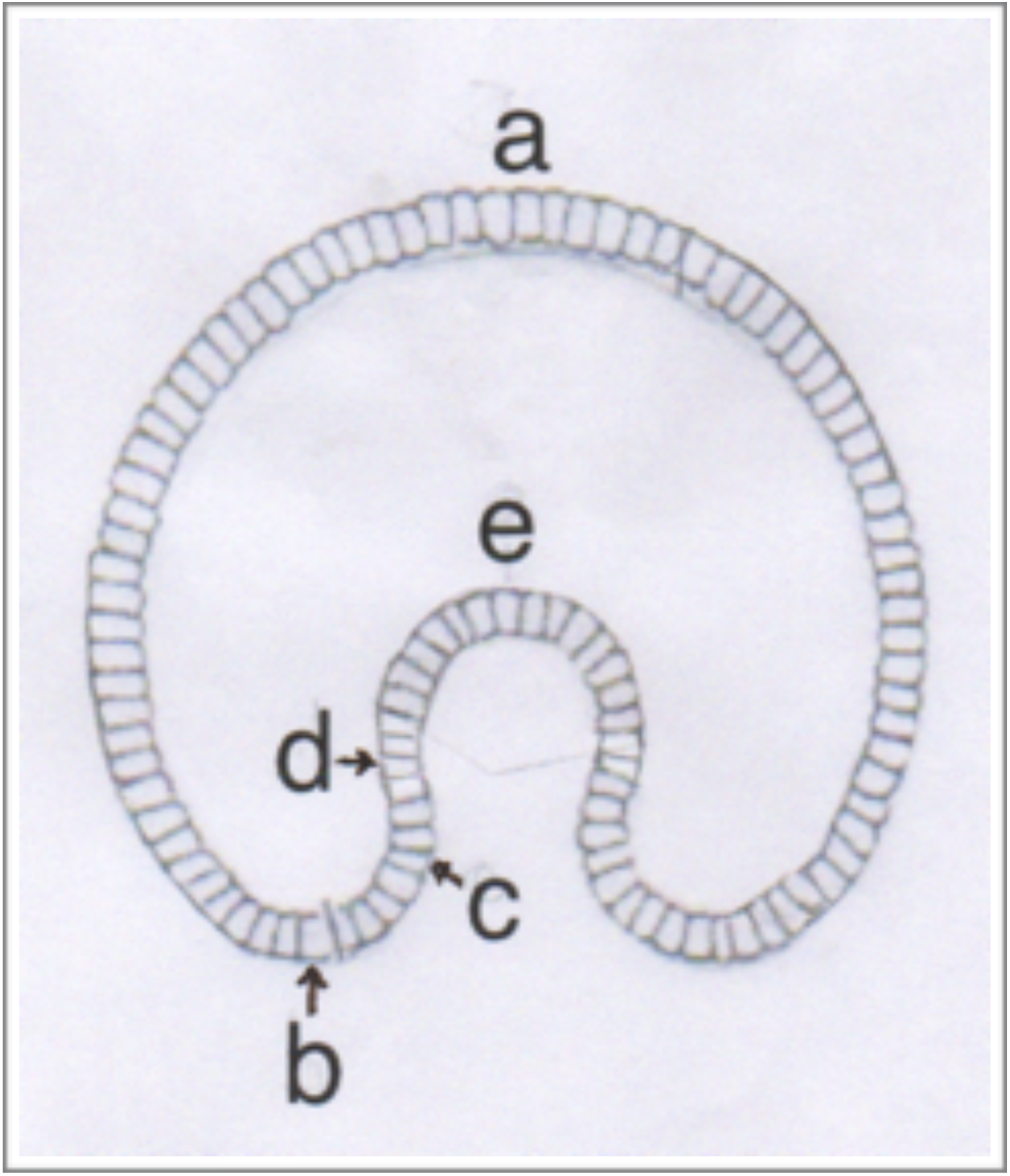
A sample coupling of patterning and epithelial shape results in a typical first-stage gastrula. The simplest assumptions are made: the thickness has been kept constant, and the dimensionless areas ‘A’ and ‘B’ have been assumed as proportional to the two co-t factors, R_1_ and R_2_. The Gauss curvature K is first zero at the tip of the gastrula ‘mouth’, at **point b**. The Mean curvature ‘H’ turns negative first at **point c,** and remains negative until the **point e**. At **point d**, the Gauss curvature K becomes zero again, and remains positive between **points d and e**. The blastopore is associated with the negative K region, between **points b and d.**

**Figure 5:**
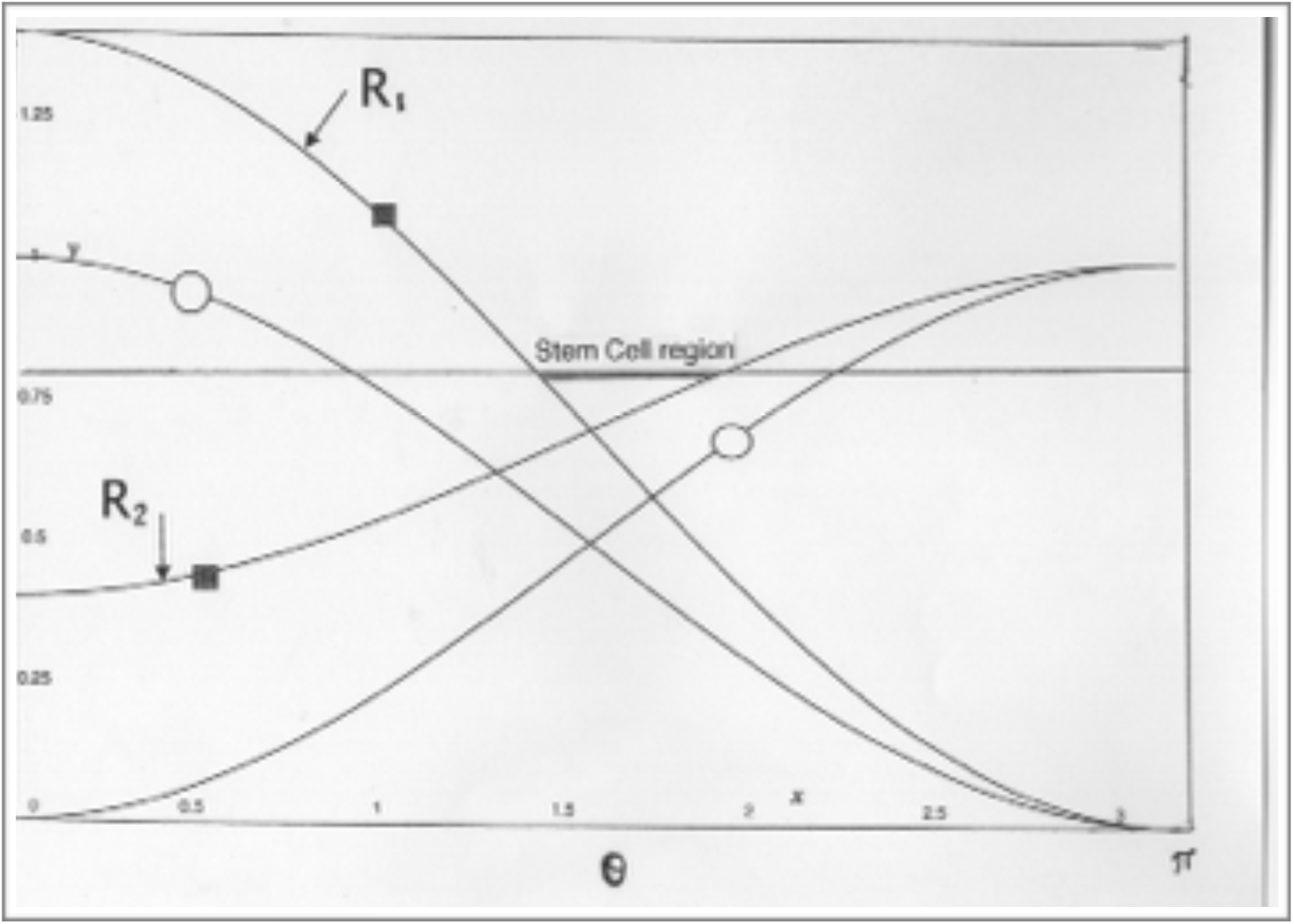
Two solution of the mathematical model are shown in the case of the small sphere (blastula) shown in Figure 3, and eqs. (4.1a), (4.1b) and (4.2). The radius is specified by eqn. (4.2) as (kR)^2^ = 2 (Appendix A). The angular region indicated by the thick horizontal line between two co-t sections is a region of no transcription; below this (here, arbitrary) defining thick line, we have A+B < 2, or K< 0, and is designated as a ‘stem cell’ region. The stem cells are controlled by the two enclosing co-t factors, R_1_ and R_2_ (here indicated by small blue squares), and have parameter values (k_1_/k)^2^ = 0.7 and (k_2_/k)^2^ = 0.3.The second two co-t regions indicated by open circles have parameter values k_1_=k_2_.

As development progresses, more complicated functional connections between A, B, h and R**_1,2_** are of course expected.

## 4) Gastrulation 1: Patterning the Blastopore and Stem Cells

The patterning of stem cells is widely discussed in the literature (c.f., PubMed for many references). Presumably they are not distributed at random in the developing organism. It is most important that a model envision the repeated generation of stem cells as the animal develops, with stem cells developing that are appropriate to a given organ or body part. For example, if a newt’s tail is excised, the tail will regenerate, by calling into action specific ‘newt-tail’ stem cells; the tail will regenerate a tail, and the remaining excised tail, if cultured, will duplicate itself. This is the general situation for excised parts, limbs, etc. (this ability, for largely unknown reasons, becomes increasingly restricted as animal complexity increases; mammals can only wound heal, and regenerate limited specified organs, and is not discussed further). Rauzi et al. (2013) have given a **review** of a number of alternate models.

Equation (A.6) for the weighted sum S of the two co-t factors given by eq. (A.7) is the famous Laplace equation and is well known to have a unique solution for any specified boundary condition. We imagine that stem cells present on any excised boundary swing into action and both regenerate and duplicate, giving the one unique solution: the weighted sum S for both the (e.g.) animal tail and the cultured excised part. A similar situation will occur for a region of the blastula; the blastula will regenerate the excised part, while the (cultured) excised part will duplicate.

The stem cells will be patterned as a a smaller region **between the two regions** of the different co-t factors (e.g., between densities of Ci (the Hh co-t) and β-catenin (the Wnt co-t)) (Nusse 2003). This can be seen by the joint solution of eqs. (A.6) and (A.8). As shown in Appendix A, the solution on a sphere of radius R, when (kR)^2^ = 2 is found from eqs. (A.6)-(A.9) to be (S = constant, and (k_1_/k)^2^ ≥ (k_2_/k)^2^),

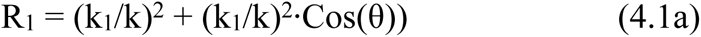

and

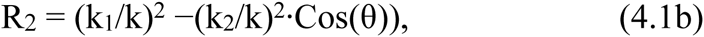

With

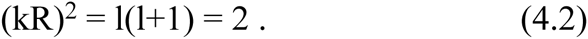

It is important to note that the area of the (spherical) blastula is given by eq. (4.2) at the start of gastrulation. It will be a large number cell diameters, and ‘k’ is provided for any specific animal by a measurement of this area ∼ R^2^.

The two co-t factors R_1,2,_ are shown on the single **small** sphere in Figure 3, for one specific but typical set of parameters, and also in Figure 5. **Also shown** are two arbitrarily chosen lower thresholds, **t1 and t_2_** in Figure 5. Figure 3 **indicates** the formation of **compartment boundaries**, the ‘M’ regions, showing two compartments for the smallest sphere, and four for the two larger spheres. A compartment, in this light, is always enclosed by a stem cell region.

A general property of the model is that each co-t factor (R_1_ and R_2_) **acts to avoid** the other, over a cellular distances of many cell diameters, of the order of k^-1^. A large enough threshold for nuclear entry of the co-t factors generally provides a region of non-transcription between the two co-t’s. This is identified as a **stem cell region** (blue in Figure 4). Such a stem cell region will be propagated repeatedly as development proceeds, necessarily providing different stem cells into the adult as required.

Figure 4 shows R_1,2_ for a typical **stage-one** gastrula. The shape will be altered somewhat by different parameter choices, but the shape of holoblastic animals, those with a small yolk, is that as largely as shown in Figure 4. Boundary conditions due to mostly impenetrable yolk (Gilbert 2012), as occurs in non-holoblastic animals such as birds, fish and reptiles, have not been considered in Figures 4 and Figure 5.

Proceeding from angle θ = 0 (point **‘a’**) to π (point **‘e’**), but left to right in Figure 4 and eq. (4.1), and from **point a to e in Figure 4,** the curvatures of the (middle) surface are specified in Figures 4 and 5 as:

**Point b** is the first place where the Gauss curvature K becomes zero, A+B= 2. Curvature K remains negative **between points b and point d**. This region of K < 0 maps out the blastopore region; in this region the cells have a ‘twist’, as indicated in Figure 1. **Point c** is where the Mean curvature ‘H’ **first becomes zero (**and A = B), and after which the apical area ‘A’ is less than basal ‘B’, the Mean curvature remains negative until **point e,** at θ = π. The **point d** is where the Gauss curvature K returns to being positive, remaining positive between **point d and point e**.

## 5. Gastrulation 2: Patterning the Blastopore and Bilaterality

The first stage of gastrulation begins during blastula growth, and the steady-state area shown in Figure 3 on the smaller sphere (left) is given by the expression (kR)^2^ = 2; (= l(l+1), where l=1). The second stage of gastrulation then proceeds, until the archenteron increases in length by about a **factor of three,** leading to a long thin tube about an area factor of three larger and **(kR)^2^ = 6,** (l=2) (Appendix B and Gilbert 2012). The area increase in the second gastrulation phase is accompanied by little growth; instead, cells largely migrate and flatten (e.g., Hardin and Cheng 1986). Figure 5 shows the two co-t functions, R_1_ and R_2,_ for two different parameter choices, as functions of the sphere angle θ. One set is shown with small black circles, while the second is the parameter choice of eq..

Bilaterality is determined during the second stage of gastrulation. This second stage results in a ‘tube within a tube’, and culminates in contact between the endodermal and ectodermal sheets, and the formation of a second opening at the point of animal/vegetal contact. In the cases that the original gastrula opening remains the mouth, the animals are known as “protostomes”, while when the second opening becomes the mouth, (still somewhat mysteriously, and not discussed further here (c.f., Martindale and Hejnol 2009)) the ‘second mouth’ animals are known as deuterostomes (e.g., vertebrates, coelenterates).

Two ‘points’ specified on opposite sides of the blastopore ring are of prime interest. A question is: how are these two important regions (often referred to as ‘organizing centers’) specified? These two (‘point-like’) regions mark the positions of activation of the ‘Brachyury’ gene, a key gene in the evolution of the mesoderm. Generally the mesoderm cells migrate between ectoderm and endoderm, one branch leading to the heart, kidney and gonads, and the second to the connective tissues, bone, muscles, tendons, and blood. The intersection of two ‘M’ regions in the blastopore ring specifies the two positions of mesodermal emergence, and is shown by the model solution in Figure 3, on the larger sphere; the mathematical details are given in Appendix B. (‘left-right’ asymmetry origins are unknown, but may arise through mechanical induction of cells; no known macroscopic aspect of nature seems to differentiate left from right; cf. Review, Levin 2005).

One ‘M’ region is shown in Figure 3 that encircles the two larger spheres from pole to pole, through the angles from θ = 0 to θ = π. This M region determines the divide **between dorsal and ventral**. This second growth phase also results in a significant lengthening of ectoderm and endoderm, accompanied by emergence of the mesoderm from the two intersection ‘points’ of the two different ‘M’ regions.

Reference to the bottom sphere of Figure 3 indicates the different unique specifications for a): the dorsal and ventral ectoderm (ab and aB), and b): the dorsal and ventral endoderm (Aℬ and Ab in Figure 3).

The conserved nature of the connections between phyla is impressive (Technau 2001). Brachyury is one of the principal transcription factors in the early determination and differentiation of mesoderm in virtually all phyla, cnidaria to vertebrates (Martin and Kimelman 2008; ibid 2012). Comparison of a variety of organisms from different phyla, from the diploblastic cnidarians to vertebrates, shows that the original sites of expression of Brachyury and its homologues are at two specified points on opposite sides of the blastopore region. From these two ‘points’ emerge the original mesodermal tissue of both of the two main animal groups, protostomes and deuterostomes.

The BMP pathway is a main signaling pathway involved in the D-V patterning of the mesoderm of the second stage gastrula of the vertebrate embryonic field, and it consists of about 10,000 cells in *Xenopus frog* embryo. BMP-4 induces the formation of epidermis and ventral mesoderm, and also blocks the induction of dorsal mesoderm, including notochord and somites, and also of dorsal ectoderm such as the nervous system (Gerhart and Kirschner 1997).

The patterns of Figure 3 are shown on a sphere for both growth phases, with no account of the shape changes. The increase in area from the smaller gastrulating sphere to the (two) larger spheres is approximately a factor of three according to the mathematics described in Appendix B. The different shapes are expected to conform closely to these patterns of Figure 3, as they remain topological spheres, until the second opening occurs, providing an anus in the case of protostomes.

## 6 Discussion and Speculation on Segments and outgrowth of limbs, antennae

One motivation for developing a universal model for patterning multicellular systems is the ubiquity of segmentation, or repeating modules in general. One persisting question involves the determination of the placement (apart from patterning) of outgrowths (or ingrowths, e.g., eyes) from the main animal body. As the animal extends its length, there is very often associated, in most phyla, a segmented body. The Helmholtz equation (2.2) gives rise to double periodicities. Figure 6 shows (two) patterns of the model on two cylinders, (viewed side on) as both cylinder length and radius increase top to bottom view, and where ‘M’ regions are indicated now simply as (black) crossing lines separating the two co-t factors.

**Figure 6:**
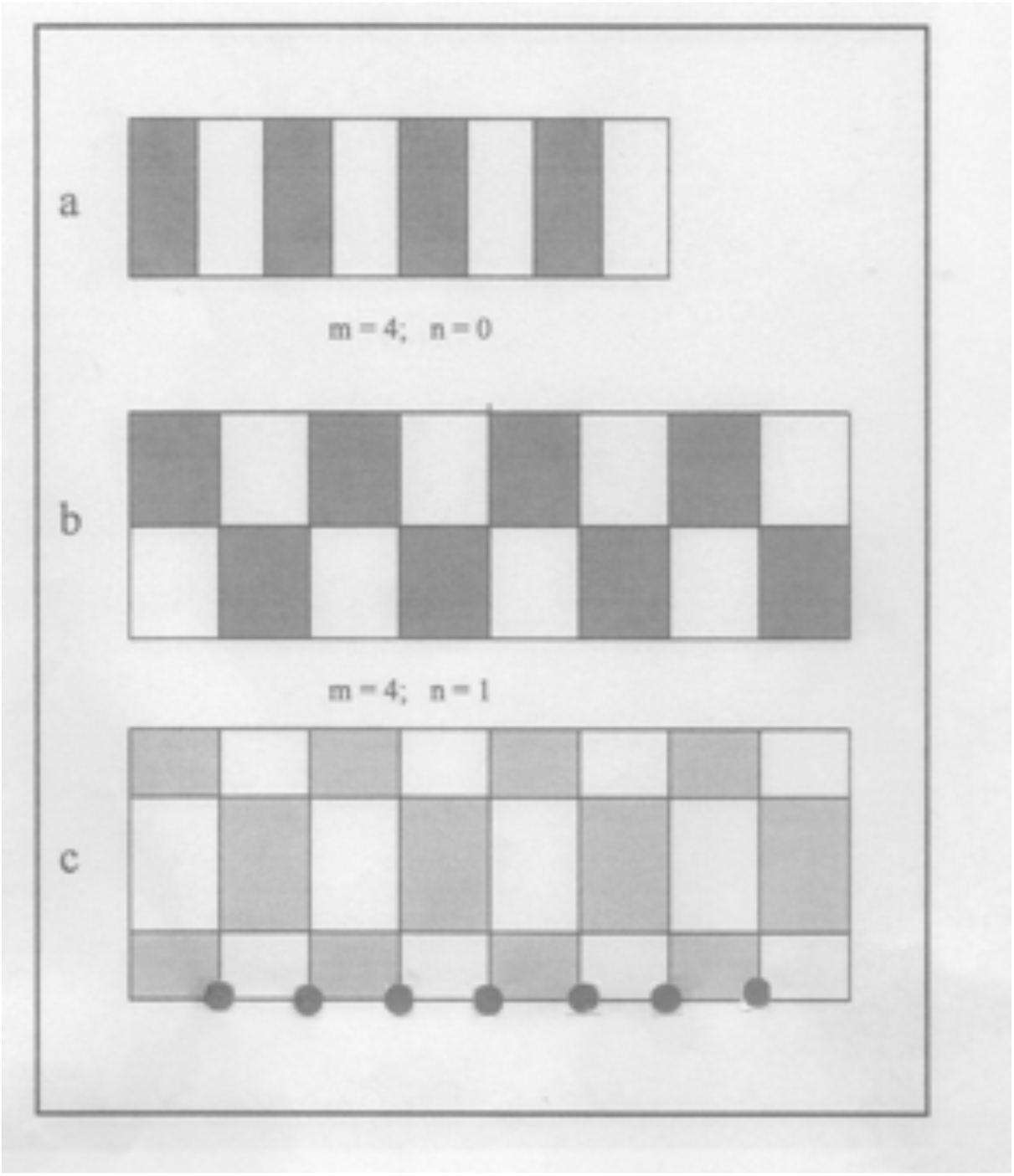
Three cylinders are shown in side view, representing the periodic patterns on a developing animal. Here the ‘M’ regions of the model are indicated only by black lines between the two co-t regions of eq. (A.8) at three indicated developmental stages, and radii. If it can be argued that limbs emerge from intersections of ‘M’ regions (as do mesodermal factors), then it is predicted that such an animal (e.g., a Centipede) will have always an odd number of leg-pairs when the HOX genes (Carroll et al., 2005) are the same for each co-t region, and shown in the bottom figure ‘c’. The head and tail regions are not shown, but under the control of other HOX genes.

One striking fact is apparent: the number of ‘M’ intersections is always an **odd number** as we go from anterior (left) to posterior (right) in Figure 6. A surprising fact about Centipedes is that, in over 3,000 different species examined, with large variations in lengths, they **always** have an odd number of leg pairs (Chipman, Arthur and Akam 2004). This is not likely adaptive, so this naturally causes speculation that the (always odd) intersections (cf. lower cylinder, Figure 6) of M regions could be the designated position of (e.g.) leg outgrowths. Carroll et al. (2005) have argued that different HOX genes along the segmented body give rise to different outcomes in animal development in general (e.g., insects with six legs, arachnids with eight). Figure 6 **(bottom)** shows only the segmented periodic part of the Centipede, i.e., the region that has activated the same HOX genes at each growth phase; Figure 6 is not showing the Centipede head and tail regions, with their own specific ‘tool-kit’ genes and ‘switches’. A ‘double segmented’ pattern (Chipman, Arthur and Akam 2004) is evident when the three regions of the present model are taken as: ‘white’ between ‘red’ and ‘blue’ (the two co-t regions the ‘red’ and ‘blue’) and separated by the white ‘M’ region. Then the periodicity along a line might be (e.g.) 1): ‘red (white) blue’; 2): ‘red (white) blue’; 3): ‘red (white) blue’, and so on. The double segment pattern assures an odd number of leg pairs. Bottom Figure 6 shows legs at the positions of the lower M intersections.

Of interest is that the crossing of two ‘M’ regions provides necessary specification of both apical/dorsal and dorsal/ventral axes at the positions of outgrowths, (e.g., legs, antennae) and ‘ingrowths’ (e.g., wing and leg discs in insects, or eye cups). The Gauss curvature ‘K’ at the M intersections are positive in both cases, K > 0, and either H > 0 (outgrowths) or H < 0 (ingrowths). Limb, etc. patterning is not considered here, only the specification of **position**, i.e., an answer to “where?”.

In the case of non-axial symmetry, most often as in plants, the present model gives rise to the usual periodic plant patterns on a cylinder (Cummings and Strickland 1998), giving the Fibonacci series as a common result. The biochemistry is totally different for plants and animals, but the basic patterning model is the same.

## 7. Speculations regarding Regeneration and Duplication

Solutions to the Laplace-Beltrami equation (2.1) are known to be unique for given boundary conditions. Here ‘S’ (for ‘weighted sum’) of the two co-t factors is the unique solution. This predicts that (e.g.) upon the bisection of the blastula, the missing piece of the larger sphere-like part will be regenerated; also, a duplicated piece of the excised part, when cultured, will also be generated. The two parts are both ‘regenerated’ from the same boundary conditions on the cut surface, resulting in two identical regenerated parts. This only occurs by a process akin to a Laplacian restorative algorithm.

The Laplace equation acts to minimize gradients, and in the time dependent version, this means that the algorithm acts to place maxima and minima of ‘S’ both on the boundary. Cell division continues until the interior gradients are sufficiently minimally smooth, and a steady state is achieved. A missing part regenerates, when supplied with nutrients, by replacing, in small increments as cells divide, each missing piece; at first by replacing missing pieces nearest to the cut surface, then replacing pieces next to this, and so on, iteratively. The new upgraded pieces at each small time interval change is the average value of its nearest neighbors. This “average value of the nearest neighbors” repeated recipe of the Laplace equation then propagates increasingly to the interior until all pieces are restored to their original values, and the completed surface **uniquely** matches the (Dirichlet) boundary conditions as demanded by Laplace-Beltrami. Cell growth has long been predicted to be sensitive to gradients. ‘Laplace-Beltrami’ viewed in the sense of: “assign the average of neighbors at each iteration” is the unique algorithm producing duplication/regeneration.

The Helmholtz equation determines a doubly periodic structure, in this case, the difference of the two co-t factors, ‘D’. It includes determination of shape and size, given boundary conditions. As mentioned earlier, the surface shape is included in the surface metric, and included in the Laplace-Beltrami operator (doCarmo, 1976). A simple one dimensional example may be helpful regarding size and periodic determination, giving boundary conditions at each end of a line: at x = 0, R_1_ = 1 and R_2_ = 0; at x = L, R_1_ = 0, and R_2_ = 1. Then the solution of the Laplace equation is simply S = C_1_, while D = C_2_·Cos(k·x), (C_2_ = C_1_k_1_^2^ = k^2^/2) and the allowed lengths are determined by kL = 2nπ, (n = ±integer). This gives the co-t factors R_1_ = Cos^2^(k·x/2), and R_2_ = Sin^2^(k·x/2), allowing different periodic solutions for different lengths ‘L’, i.e., providing the periodic extensions in time as happens with very many phyla with growth.

## 8. Summary, Conclusions and Speculations

We have taken the view that embryonic animal development consists of two main inextricable entities, and the interplay between two interlocking parts. The one, and the most developed here, is that of the interaction of two signaling pathways, this providing the ‘patterning’ at each growth cycle. The ‘patterning’ of compartments and stem cells requires only short range diffusion, but nevertheless results in long range patterning (∼ k, an inverse length, cf. eqn. (2.3)).

Perhaps equally important second aspect is the interaction of this ‘patterning’ with the ubiquitous fiber mesh, or fiber network. The two, fiber network and pattern, are inextricably connected; each regulates the other. Fiber networks are numerous and diverse, from extracellular cell fibers to nerve fibers. The only fiber interaction discussed here is that of Wnt and Hh with the three key fibers of the cytoskeletal network.

The extracellular matrix (ECM) is a collection of extracellular molecules secreted by cells that provides structural and biochemical support to the surrounding cells. Because multicellularity evolved independently in different multicellular lineages, the composition of ECM varies between multicellular animals (and plants); cell adhesion, cell-to-cell communication and differentiation are common functions of the ECM.

A compression buffer against the stress placed on the ECM is provided by polysaccharides and fibrous proteins that fill the interstitial space. Basement membranes are sheet-like depositions of ECM on which various epithelial cells rest. The ECM serves many functions, providing support, segregation of tissues, and regulation of intercellular communication. Collagens, the most abundant protein in the human body, are the most abundant protein in the ECM. It accounts for 90% of protein content of the bone matrix. Fibrillar proteins give structural support to resident cells. Collagen is exocytosed in precursor form (procollagen), where it is then cleaved by procollagen proteases to allow extracellular assembly.

Elastins, in contrast to collagens, give elasticity to tissues, allowing them to stretch when required, and then return to their original shape. Blood vessels, the lungs, skin and other tissues contain high amounts of elastins. Elastins are synthesized by fibroblasts and smooth muscle cells. Disorders such as Williams syndrome are associated with deficient or absent elastin fibers in the ECM.

Changes in cell morphology, intracellular transport and exocytosis all depend on the activities of cytoskeletal filamentous structures – microtubules, microfilaments and intermediate filaments. Different regulatory pathways regulate cytoskeletal functions, anything involving cell shape changes, such as seen in Section 3.

The view adopted in this work was motivated partly by the observation, (and the assumed extension to several or many other SPs), that both Wnt and Hh, so ubiquitous in embryogenesis, and both have (at least) two modes of operation: one mode of each is directed toward gene transcription, e.g., β-catenin accumulates around the nucleus in the case of Wnt. The second mode, acting when (e.g., β-catenin level is low in the case of Wnt), Wnt, Hh, BMP, Notch, TGF-β, etc. are recruited in various fiber regulation requirements. Wnt and Hh are known regulators of the three fibers of the cytoskeleton.

The above work presents a possible, and hopefully plausible, model that correlates a few of the most common aspects of the earliest embryological development of animals. It is hoped to provide some insight into the emergence of animal from their single cell progenitors some 600 my ago. A notable aspect of the present model is the separation of each compartment into two regions, each extending over many cell diameters, (∼ k^-1^) at each ‘growth cycle’ of each previous compartment. Compartment boundaries are specified as the separations between co-t factors, delineated by the ‘M’ regions. The separation between the two regions (called ‘co-t’ regions, co-transcription regions) is identified as regions of stem cells for this specific cycle, an intermediate region where transcription of this cycle does not occur.

The first phase of gastrulation includes formation of the the blastopore, as well as the two places on opposite sides of the blastopore marking the emergence of the mesoderm. The second phase of gastrulation, accompanied by specification of the dorsal/ventral axis is given by the model. The ‘double segmental’ nature of the model provides a model for the very notable **‘odd-leg pair’** number character of the numerous species (>3,000) of Centipede.

After the emergence of the mesoderm, complexity increases at a rapid rate. The profusion of signaling pathways now being studied is impressive (Berridge 2014). One of the most conspicuously absent universal from the present model are nerve nets. Jellyfish, Hydra and starfish have intricate nerve nets, and worms distinguish light from dark. Nerve cells in crayfish are basically similar to vertebrate (or human) nerve cells (Sacks 2014). The emergence of the mesodermal cells at the time of the dorsal/ventral determination provides great diversity, and is a subject not touched on here.

## Appendix A: Mathematical Formulation of the Model

The model of the previous section is now most simply expressed mathematically. In the first instance, each ligand density in the extracellular space changes due to two reasons: first it can diffuse over short distances between the cells, and second, it is increased by the density of intracellular co-transcription (‘co-t’) factor. This latter increase of one type of secreted extracellular ligand is attenuated in turn by the presence of the second intracellular co-t factor.

This is expressed mathematically by the two equations (where L_1,2_ denotes either L_1_ **or** L_2_, etc.),

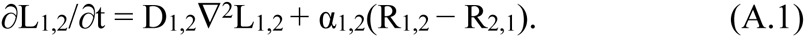

The operator ∇2 is known as the ‘Laplacian’(e.g., Morse and Feshbach 1961), or more exactly as the Laplace-Beltrami operator, and ∇2 L represents the difference between specific ligand ‘L’ in a given small region minus the average of the same ligand density of the nearest neighbors, thus connecting cell to neighboring cell. The ‘Laplace-Beltrami’ operator also importantly involves the epithelial shape.

The time change of each ‘co-t’ factor, R_1_ or R_2_, is given most simply by the two equations

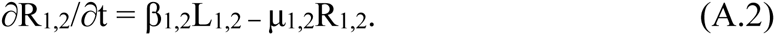

Together with an **important lower limit on the density of R_1,2_** needed for gene activation, these four equations, (A.1) and (A.2) constitute the model. The upper limit for R_1,2_ is provided of course by the number of cell surface receptors of each kind.

Of most interest is the steady state version of the model, when the left hand sides of eqs. (A.1) and (A.2) are zero, that is, ∂/∂t = 0. As will be seen, the dynamics is such that the two quantities R_1_ and R_2_ act as time increases so as to increasingly avoid each other as the two increase in density, while at last leaving a (usually thin) region between, labelled the ‘M’ (or middle) region. The size of ‘M’ is determined by the lower threshold for allowing entry of the various co-t factors into the nucleus (two factors per growth phase), where they cooperate with nuclear partners and with the DNA and switches to transcribe a given gene or gene network. The lower limit provides an important role for the nucleus in the patterning.

The steady state equations are at once, from eq. (A.2)

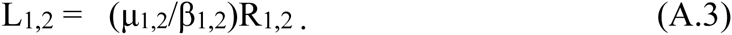

Now inserting eq. (A.3) into the steady state version of eqs. (A.1) we obtain the two equations (A.4), where

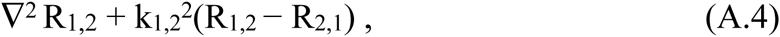

where

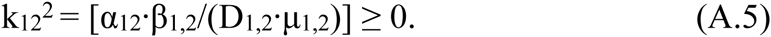

The two equations (A.4) can be added and subtracted, giving two key equations of the model, eqs. (A.6) and (A.8), the **sum ‘S’ and difference ‘D’** defined by

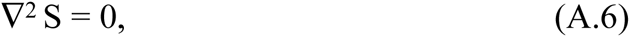

where

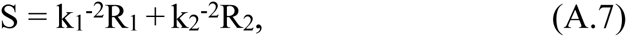

and

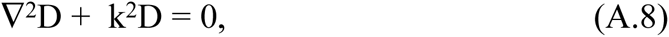

where

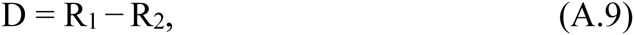

and 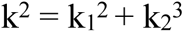.

Equations (A.6) and (A.8) are the two steady state equations of the model. The simplest solution on a sphere of eq. (A.8) is D = const·P_1_^0^=const·Cos(θ), and from eqs. (A.8) and (A.6) together it follows generally that R_1_ = (k_1_/k)^2^S + (k_1_^2^/k^2^), and R_2_ = (k_1_k_2_/k)^2^S - (k_2_^2^/k^2^)D, where S and D are solutions to the eqns (A.6) and (A.8), then

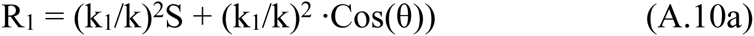

and

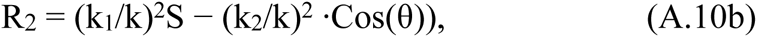

and are eq. (4.1) of the text, (here we take S = 1= constant), and two examples are shown in Figure 5. The constant ‘S’ is ∼ the maximum density of co-transcription factors R_1,2_ in the cells under consideration. In the case that k_1_ = k_2_, then R_1,2_ equals Cos^2^(θ/2) and Sin^2^(θ/2).

## Appendix B

The four regions of the **second growth phase** of the gastrulating embryo, shown on the larger sphere at the right of the Figure 3, are given by the model eqs. (A.6) and (A.8). The conditions that R_1_ and R_2_ be positive gives the solution

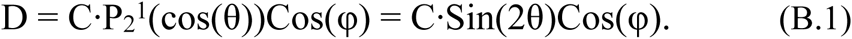

Here P_2_^1^ is an associated Legendre function, and C is a constant.

The co-t factors R_1,2_ are then obtained from a combination of eqs. (A.7) and (A.9) to give, taking S = 2C/k^2^, and

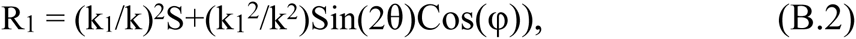

and

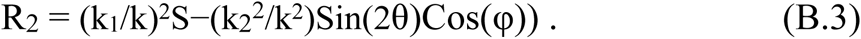

Equations (B.2) and (B.3) shown on a sphere in Figure 3 (top right) as ‘red’ and ‘blue’ regions; ‘S’ is a constant, again taken as unity, determining the the maximum of R_1,2_ when θ = 0 or θ = π.

The corresponding **sphere radius** is now given by (kR)^2^ **= 6,** (with l = 2, and l(l+1) = 6), and is a **factor of three increase** in area over the first growth phase of Appendix A (cf. S. Gilbert, 2012). One of the two ‘M’ regions encircles the sphere from pole to pole, passing through the angles θ = 0 and θ = π. This M region determines the divide between **dorsal and ventral**.

This second growth phase results in a significant lengthening of ectoderm and endoderm, accompanied by emergence of the mesoderm from the **two intersection ‘points’** of the two different ‘M’ regions shown in Figure 3 as a projection onto the two larger spheres.

Reference to the bottom sphere of Figure 3 indicates the different unique combined gene specifications corresponding to **a**): the dorsal and ventral ectoderm (ab and aB) and **b**): the dorsal and ventral endoderm (Aℬ and Ab).

The patterns of Figure 3 are shown on different size spheres for the two growth phases, with no account taken of the gastrulation shape changes. The gastrula shapes are expected to conform closely to these patterns, and they remain topological spheres until the second opening forms, when the Gauss integral of K over all area changes from 4π to zero.

## Acknowledgement

A critical reading of the manuscript by M. Tavis is appreciated.

